# Mathematical Modelling of Stochastic Gene Expression Quantifies mRNA Heterogeneity in Cancer Cell Populations with Extrachromosomal DNA

**DOI:** 10.64898/2026.10.07.757305

**Authors:** Poulami Somanya Ganguly, Weini Huang

## Abstract

Oncogene amplification on circular extrachromosomal DNA (ecDNA) has been linked to poor prognosis and higher treatment resistance in multiple types of human cancer. ecDNA are mobile genetic elements lacking centromeres that are partitioned unevenly into daughter cells at mitosis. While random segregation of ecDNA contributes to gene-copy-number heterogeneity among tumour cells, how ecDNA contribute to phenotypic heterogeneity, via mRNA and proteins, and how this affects targeted therapy outcomes is still not understood. In fact, cancer cell populations with ecDNA show remarkable heterogeneity in mRNA concentrations, which is not explained by gene-copy-number heterogeneity alone. In this paper, we use a stochastic model of gene transcription to mathematically model mRNA production and decay in cancer cells with ecDNA, extending a method that has been used to study phenotypic heterogeneity in eurokaryotic and prokaryotic cells. Using our model, we explore the space of possible mRNA distributions from a given ecDNA distribution, under different model parameter regimes. Our model is able to produce realistic mRNA distributions and we use it to estimate transcriptional parameters from single-cell measurements of ecDNA and mRNA counts in cell-line data. We discover a remarkable similarity in estimated parameters across cell lines and amplification types, indicating similar processes underlying transcription in ecDNA and chromosomal amplifications. We also model genotoxic therapy and use our model to predict the mRNA composition of persisting cells, showing the usefulness of such a modelling framework to understanding the origins of treatment resistance in ecDNA-containing cancers.

## 1 Introduction

Extrachromosomal DNA (ecDNA) are circular particles of genetic material in the cell nucleus that have been observed to carry important oncogenes in several types of human cancer. The presence of ecDNA has been linked to aggressive tumour growth, therapy resistance and poor patient survival. An important characteristic of ecDNA that distinguishes it from chromosomal DNA is that ecDNA lacks centromeres and, as a result, is partitioned unequally between daughter cells during cell division. Consequently, oncogene amplifications on ecDNA show greater copy-number variability within a population than chromosomal amplifications, contributing to faster tumour evolution [1].

In previous studies, gene (or ecDNA) copy number has been taken as a proxy for the phenotypic status of cells such as fitness [1, 2]. However, experimental data suggests that ecDNA copy number alone may not always predict cell phenotype, which is better understood from the mRNA or protein content of the cell. Nathanson et al reported that glioblastoma cells with low levels of EGFRvIII protein could still harbour large copy numbers of ecDNA carrying the mutant *EGFR* gene in treatment-naive conditions [3]. In single-cell data too, neuroblastoma cells with similar ecDNA numbers and amplification were observed to vary in their mRNA content [4]. These studies indicate that mRNA production from ecDNA is itself heterogeneous and must be understood quantitatively as the next step for a more accurate picture of ecDNA-driven heterogeneity. Moreover, gene expression heterogeneity, which does not require changes in gene copy number, might be an additional mechanism of modulating phenotypic status of cells. To quantitatively understand this heterogeneity, we model transcription on ecDNA as a stochastic process, using a framework that has been used to study gene expression heterogeneity in both eukaryotic and prokaryotic cells.

Noisy gene expression leads to phenotypic differences in isogenic populations, for example in bacteria where this variability has been shown to promote survival in fluctuating environments [5, 6]. Gene expression noise due to small molecule numbers is also observed in eukaryotic and mammalian cells. In the latter, transcription proceeds in bursts rather than continuously, leading to an mRNA distribution that is non-Poisson. The simplest mathematical model that recapitulates such a distribution consists of two gene states: active (transcribing) and inactive (non-transcribing) with random switching between the two. This two-state model of transcription has been used to explain transcriptional bursting, bacterial persisters and even the appearance of coordinated high-expression states in cancer cells [7, 8, 9].

Unlike in the studies above, in ecDNA-containing cancer cells, we observe two interacting sources of heterogeneity: gene copy number and transcriptional stochasticity. Thus, it is important to have a framework to study the effect of stochastic gene expression while extending existing models to include copy-number variability. While it is possible to analytically solve for steady-state mRNA distributions when cells have only one gene copy, our case is analytically intractable and must be solved numerically through stochastic simulations. We are able to generate putative mRNA distributions in our simulations and compare them against experimental data to estimate gene switching parameters. Our model is able to match experimental mRNA distributions in certain parameter regimes. We obtain different values of switching parameters for genes on the same amplicon. Distinct parameter regimes are obtained for different genes carried in the same amplification contexts. Moreover, for the limited genes shared among different neuroblastoma cell lines, we observe remarkable similarity in terms of parameter regimes for the same gene in different amplification contexts and cell lines.

Our mechanistic model provides a framework with which to quantify gene expression across genes and amplification types, and is a first mathematical step towards understanding transcriptional heterogeneity in cells carrying ecDNA. In addition, we are able to use our model to predict the mRNA composition of cells that survive genotoxic therapy, which is commonly used in neuroblastoma. Our predictions indicate that the fraction of cells with high mRNA and low ecDNA content, which are phenotypically active but not amenable to further genotoxic therapy, varies depending on the amplified oncogene, suggesting a diversity that might underlie relapse.

The paper is structured as follows. In Section 2, we provide details of our mathematical model and simulation framework as well as a sketch of our parameter estimation method. In Section 3, we demonstrate our results, starting from an exploration of the different mRNA distrbutions that can arise from the same ecDNA distribution to parameter estimates from experimental data and predictions after genotoxic therapy. We conclude with a brief discussion of the future applications and extensions of our approach in Section 4.

## 2 Methods

### 2.1 Stochastic model of gene transcription in cells with ecDNA

We extend a two-step transcriptional model to incorporate the impact of underlying heterogeneity in the number of gene copies among cells. This captures for example the scenario in cancer cells with extrachromosomal amplifications or homogeneously staining regions (HSRs), where the copy number of genomic structures is numerous and variable across cells in a population. The two-step transcriptional model can be summarised in the following set of reactions:

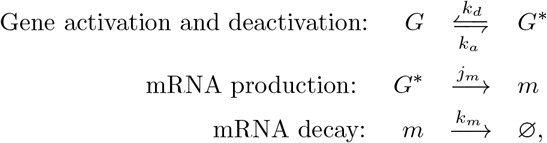

where *G* represents the inactive gene, *G*\* the active gene, *m* the mRNA and ∅ a neutral decay product. The rates of gene activation, gene deactivation, mRNA production per activated gene copy and mRNA decay are *k*_*a*_, *k*_*d*_, *j*_*m*_ and *k*_*m*_, respectively.

For cells with a single gene copy, this model has been analytically solved in the limit of large *j*_*m*_, where mRNA production is deterministic when a gene copy is in the activated state [8]. We use the same limit to obtain numerical solutions for our system with intratumour heterogeneity in DNA copy numbers. The two relevant parameters in this regime are the rescaled activation rate *r*_*a*_ = *k*_*a*_*/k*_*m*_ and the rescaled deactivation rate *r*_*d*_ = *k*_*d*_*/k*_*m*_. In the supplementary materials, we show analytical results for the special case of two gene copies. For the more general case such as in ecDNA and HSRs, where cells can have multiple and various gene copies, we simulate this system of reactions using the Gillespie algorithm and obtain the DNA and mRNA copy number distributions [10]. A schematic of our framework is shown in Figure 1 using ecDNA as an example, where we focus on the simplest scenario assuming independent stochastic switching of individual genes.

**Figure 1.**
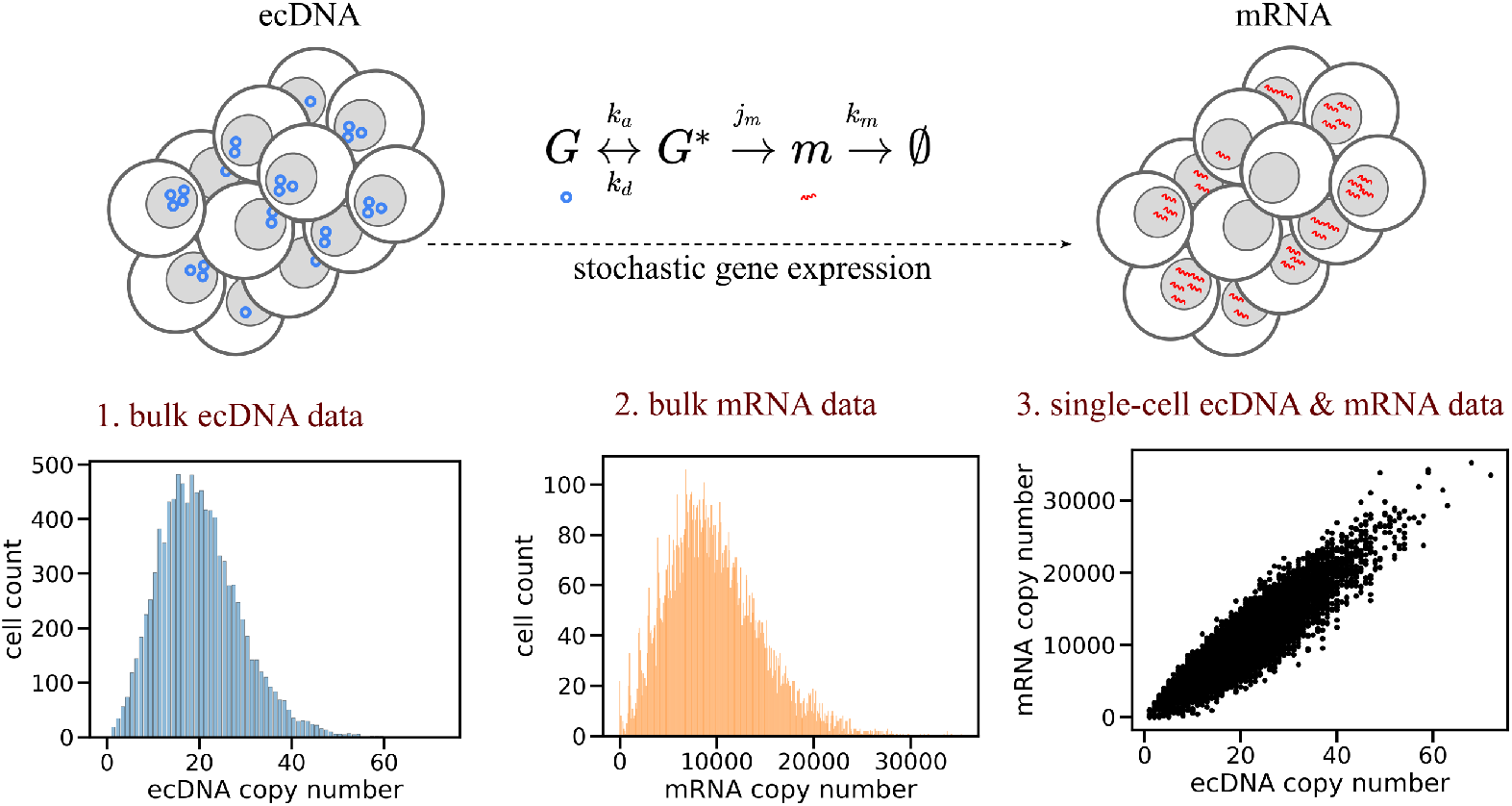
Schematic of modelling approach. A population of cells with varying copy numbers of ecDNA is taken as the starting point. Stochastic gene switching on ecDNA (blue circles) leading to mRNA production (red wavy lines) and decay is simulated to yield mRNA copy numbers. Three kinds of relevant biological data are extracted from the simulations: 1. bulk ecDNA data, 2. bulk mRNA data and 3. single-cell ecDNA and mRNA data. The last corresponds to the joint distribution of ecDNA and mRNA copy numbers while the first two correspond to the marginals.

We extract ecDNA and mRNA copy numbers in each cell at steady state from our simulations and obtain three kinds of distributions (Figure 1). The marginal distributions, namely, the distribution of mRNA copy numbers (shown in orange) and the distribution of ecDNA copy numbers (shown in blue) give us bulk information. Single-cell information can be obtained from the joint distribution of DNA and RNA copy numbers. We then use these distributions to estimate model parameters from experimental data.

### 2.2 Estimation of model parameters from data

We simulate mRNA copy numbers from cellular DNA copy numbers for various values of the model parameters, *r*_*a*_ and *r*_*d*_ (Table 1). Lower parameter values indicate a regime of slow gene switching – where stochastic switching of the gene occurs at longer time scales compared with the decay of mRNA – whereas high parameter values correspond to fast gene switching. The relative magnitude of *r*_*a*_ and *r*_*d*_ indicates the state the gene primarily exists in; for instance, *r*_*a*_ *> r*_*d*_ indicates that the gene is more likely to be active, and thus transcribing, than inactive during the approach to steady state.

**Table 1.**
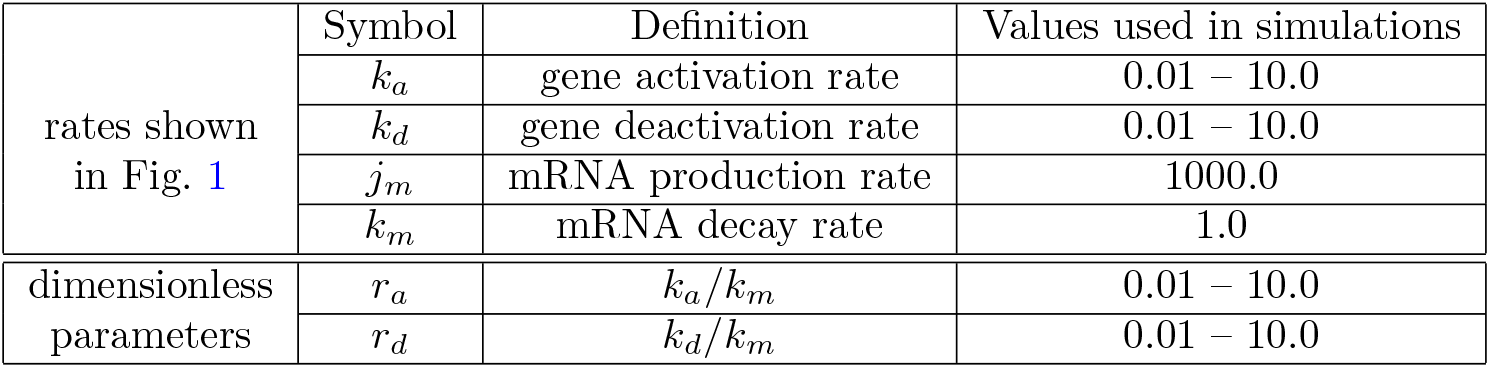
Rates in stochastic gene expression model.

To estimate the values of these model parameters from data, we use the following protocol, which is based on an approximate Bayesian computation (ABC) estimation of parameter values in evolutionary dynamics [11]. First, using experimentally measured ecDNA copy numbers we simulate a range of mRNA counts for different values of *r*_*a*_ and *r*_*d*_. Next, we use correlated single-cell mRNA measurements to compute the Wasserstein distance between the true and simulated mRNA distributions, denoted by *m*_true_ and *m*_sim_(*r*_*a*_, *r*_*d*_) respectively. For each parameter set, the Wasserstein distance is defined as

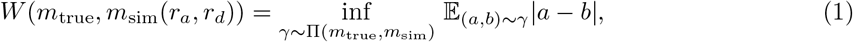

where Π(*m*_true_, *m*_sim_) indicates the set of joint probability distributions whose marginals are *m*_true_ and *m*_sim_(*r*_*a*_, *r*_*d*_), and *a* and *b* are samples from the joint distribution *γ*. Finally, parameter values that correspond to the closest match are selected based on a selection criterion. In our case, we choose to retain the parameter values for which the Wasserstein distance between true and simulated mRNA distributions lies in the lowest 10% of all distances obtained for a given gene over all random realisations of the simulation. This procedure corresponds to taking a uniform distribution of parameter values as a prior distribution and obtaining a posterior distribution adjusted according to experimental data.

We can also estimate model parameters from the marginal distributions, without using the full information in the joint distribution (Supplementary Figures S3 and S4). To do this, we bin the simulated and true marginal distributions and compute the Wasserstein distance between the binned distributions. The number of bins is an additional free parameter in the fitting procedure. Our results showed that posterior distributions obtained using the joint distribution and those obtained using only the marginal distributions are similar to each other for different bin sizes (Supplementary Figure S3). Posterior distributions obtained using the joint distribution are narrower. In order to take into account the full information present in matched single-cell DNA-mRNA data, we will present parameter estimation results using the joint distribution in the following sections.

## 3 Results

### 3.1 Transcriptional stochasticity leads to different mRNA distributions for the same ecDNA distribution

Given the same underlying ecDNA copy number distribution, we observed different mRNA copy number distributions due to transciptional stochasticity, depending on the gene activation and deactivation rates, *r*_*a*_ and *r*_*d*_. We divided our parameters into four representative regions: (i) *r*_*a*_ *< r*_*d*_, (ii) *r*_*a*_ = *r*_*d*_, (iii) *r*_*a*_ *> r*_*d*_ and (iv) *r*_*a*_ ≫ *r*_*d*_, and observed a gradient change both in the joint mRNA–ecDNA distribution (lower panels in Figure 2) and in the mRNA marginal distribution (orange in upper panels) along those parameter regions. In most parameter regions (first three panels), there is a single peak in the mRNA copy number distribution, whose position with respect to the peak of the DNA distribution varied according to the relative values of *r*_*a*_ and *r*_*d*_. When *r*_*a*_ *< r*_*d*_, the mRNA distribution has a relatively distinct shape from the underlying ecDNA distribution, which is very narrow and peaked at a relatively low mRNA copy number. We shall demonstrate later that this agrees with patterns observed in genes with low per-copy expression like *MYCN* [4]. When *r*_*a*_ = *r*_*d*_, on average half of the ecDNA copies were transcribed. When *r*_*a*_ *> r*_*d*_, the single-peaked mRNA distributions became even wider and coincided with the corresponding ecDNA distribution. This means that the shapes of mRNA and ecDNA distributions are similar and can be scaled by a linear factor. This corresponds to the regime when all ecDNA are active and transcribing, thus the lowest amount of gene switching stochasticity.

**Figure 2.**
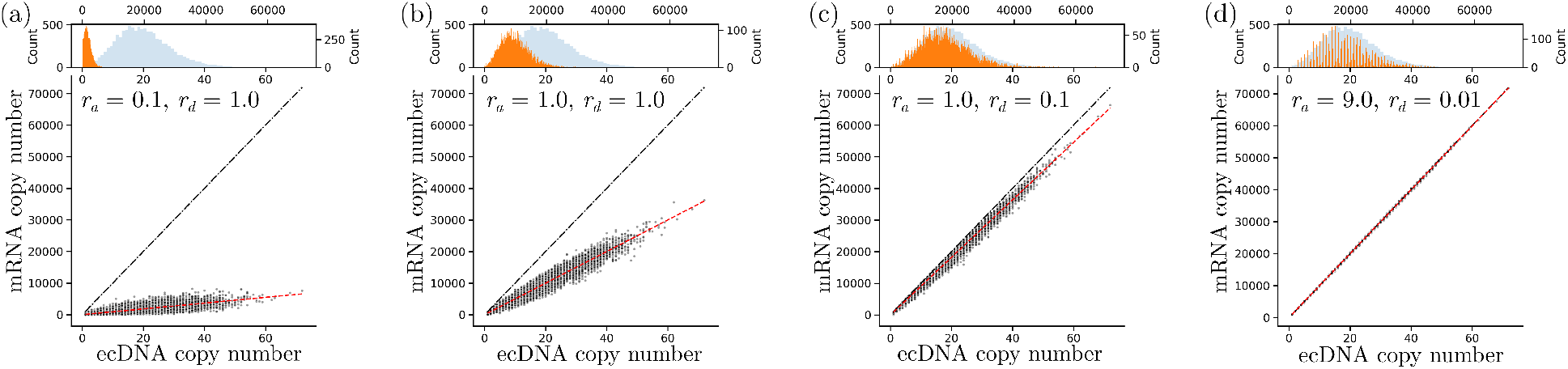
Joint distributions of ecDNA and mRNA copy numbers for different values of *r*_*a*_ and *r*_*d*_ with marginal distributions shown above. The dashed-dot and dotted lines in each plot denote the expected behaviour when all gene copies are active and when half of the total number of gene copies is active, respectively. The red line shows the linear regression fit for each set of parameter values.

In the joint distribution, we observed an increase in the slope of the best-fit line (shown in red) with increasing *r*_*a*_, and a decrease in the spread of the distribution, which corresponds to the variance in mRNA copy numbers among cells with similar ecDNA copy numbers. The slope of the best-fit line corresponds to the average effective per-copy expression, which coincides nearly with the expected value if all ecDNA copies are active (the slope of the black dashed-dot line) when *r*_*a*_ *> r*_*d*_ (Figure 2c). In contrast, when *r*_*a*_ = *r*_*d*_, at steady state half of all ecDNA are active leading to the slope of the red line lying below that of the black line. As expected, the slope is lowest when *r*_*a*_ *< r*_*d*_ (Figure 2b,a).

Interestingly, we observe multimodality when *r*_*d*_ is below a threshold value, approximately smaller than 1*/n*, where *n* is the average DNA copy number per cell. In this parameter region, multiple peaks exist in the mRNA copy number distribution separated by areas where the expression level is negligible (marginal distribution shown in orange in Figure 2d). In the Supplementary Materials we show analytically that for *n* = 2 and *r*_*a*_ = *r*_*d*_, the threshold for multimodality is 1*/*2, and conjecture that in general this occurs when *r*_*a*_ and *r*_*d*_ are less than 1*/n*.

### 3.2 Different gene switching rates provide quantitative fits to cell-line data at single-cell resolution

We applied our model to publicly available single-cell data in neuroblastoma cell lines containing ecDNA or HSR [4]. Figure 3 shows best fit results (red dots in upper panels) and posterior distributions for *r*_*a*_ and *r*_*d*_ (lower panels) for the cell line CHP212. All genes in this cell line are observed to be amplified on ecDNA and are present in the MYCN amplicon. The joint distribution of ecDNA and mRNA copy numbers (black dots in left panels) varies across the different genes, with *DDX1* being more highly expressed compared to *MYCN*, for example. Our best fits capture the observed experimental distributions well. Different values of *r*_*a*_ and *r*_*d*_ are estimated for oncogenes present in the same amplicon, which holds for other cell lines (see Supplementary Figures S1 and S2). In CHIP212, for the genes *GREB1* and *NTSR2*, which are only negligibly expressed, we obtained parameter fits in the region of *r*_*a*_ ≪ *r*_*d*_ (Figure 3a,b). Similarly, for *MYCN* we obtained a fit for *r*_*a*_ that is nearly two orders of magnitude smaller than *r*_*d*_ (Figure 3c). For *TRIB2, LPIN1* and *DDX1*, the most highly expressed genes in this cell line, we obtained smaller values of *r*_*d*_, which are much closer to the fitted *r*_*a*_ values (Figure 3d–f).

**Figure 3.**
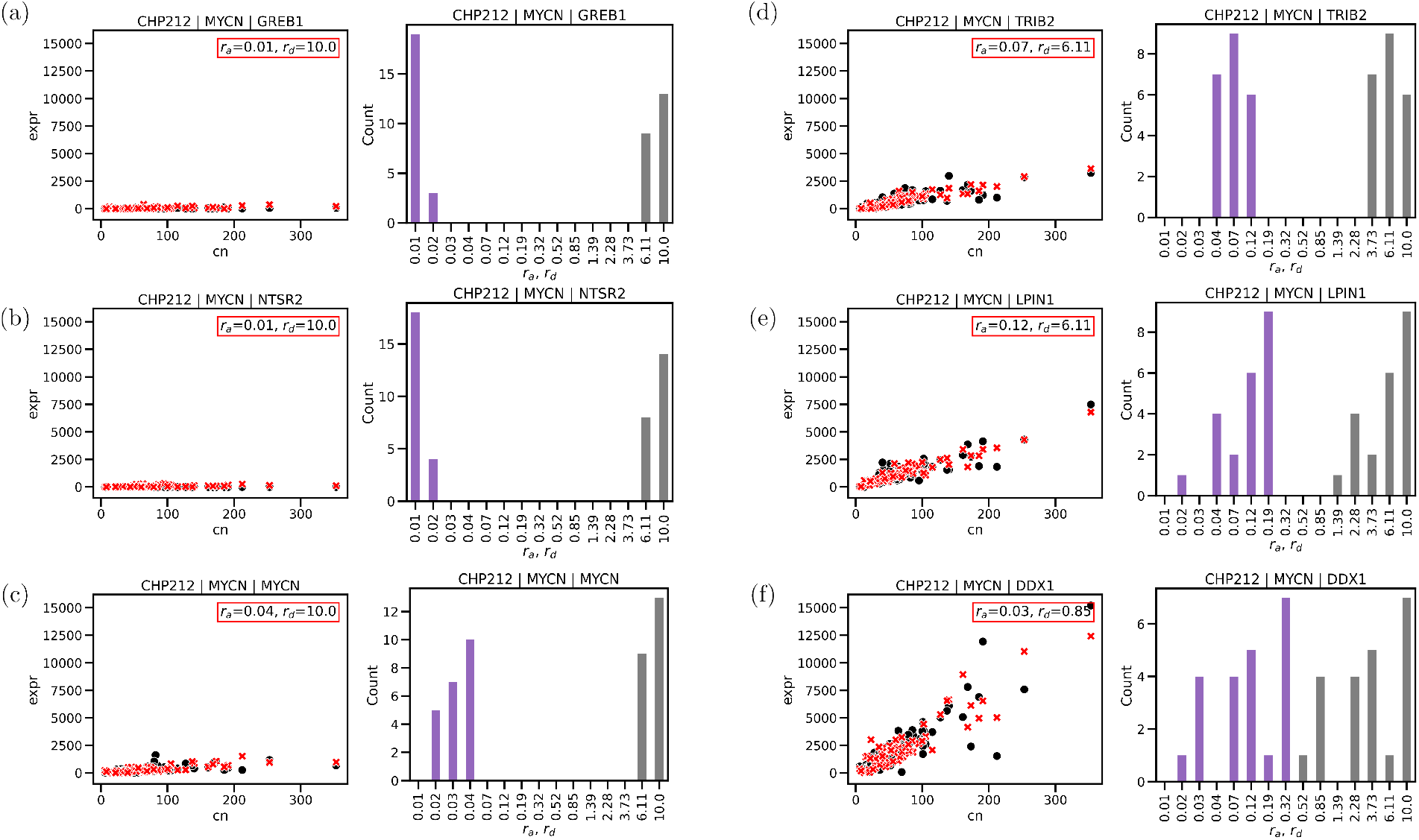
Best fit of switching parameters to genes on the same amplicon (MYCN) on the same cell line (scatter plots) and posterior distributions (histogram plots) of accepted parameter values (*r*_*a*_ in purple and *r*_*d*_ in grey)

The gene *MYCN* is highly expressed in developing tissue but downregulated thereafter [12]. The amplification of *MYCN* is widely accepted as a marker for high-risk neuroblastoma [13]. Our estimation of a low activation rate and a high deactivation rate for *MYCN* agrees well with the fact that high copy numbers of *MYCN* are often observed in patients [14]. A combination of low activation and high deactivation leads to low effective expression per copy number (as measured by the slope of the linear fit), which is likely compensated for by high ecDNA copy numbers. See the Supplementary Figures S1 and S2 for additional fits of *r*_*a*_ and *r*_*d*_ in three other cell lines which lead to similar observations. See Supplementary Figure S3 for posterior distributions obtained using bulk mRNA data.

### 3.3 Switching parameters are conserved across different amplicons and cell lines

While there are only two oncogenes, *MYCN* and *DDX1*, shared among ecDNA and HSR cell lines, we observed remarkable similarity in the activation and deactivation rates, *r*_*a*_ and *r*_*d*_, estimated for these two genes across different amplicons and cell lines. Figure 4 shows posterior distributions for the genes *MYCN* and *DDX1* in all four cell lines: CHP212 and TR14 with ecDNA amplification, and IMR5/75 and Kelly with HSRs. Across all four cell lines, *MYCN* expression corresponds to a region where *r*_*a*_ ≪ *r*_*d*_ and *DDX1* corresponds to a regime where *r*_*a*_ ≈ *r*_*d*_. Interestingly, we obtain the same pattern when estimating switching parameters from bulk mRNA data (Figure S4). Further validation with more oncogenes in different cell lines or clinical data is required to ascertain whether the observed similarity between ecDNA and HSR is a general pattern. However, our model provides a quantitative tool to identify underlying similarities or differences in ecDNA–mRNA relationships of the same oncogenes amplified under different contexts, although the absolute distributions of ecDNA and mRNA copy numbers vary between those cell lines.

**Figure 4.**
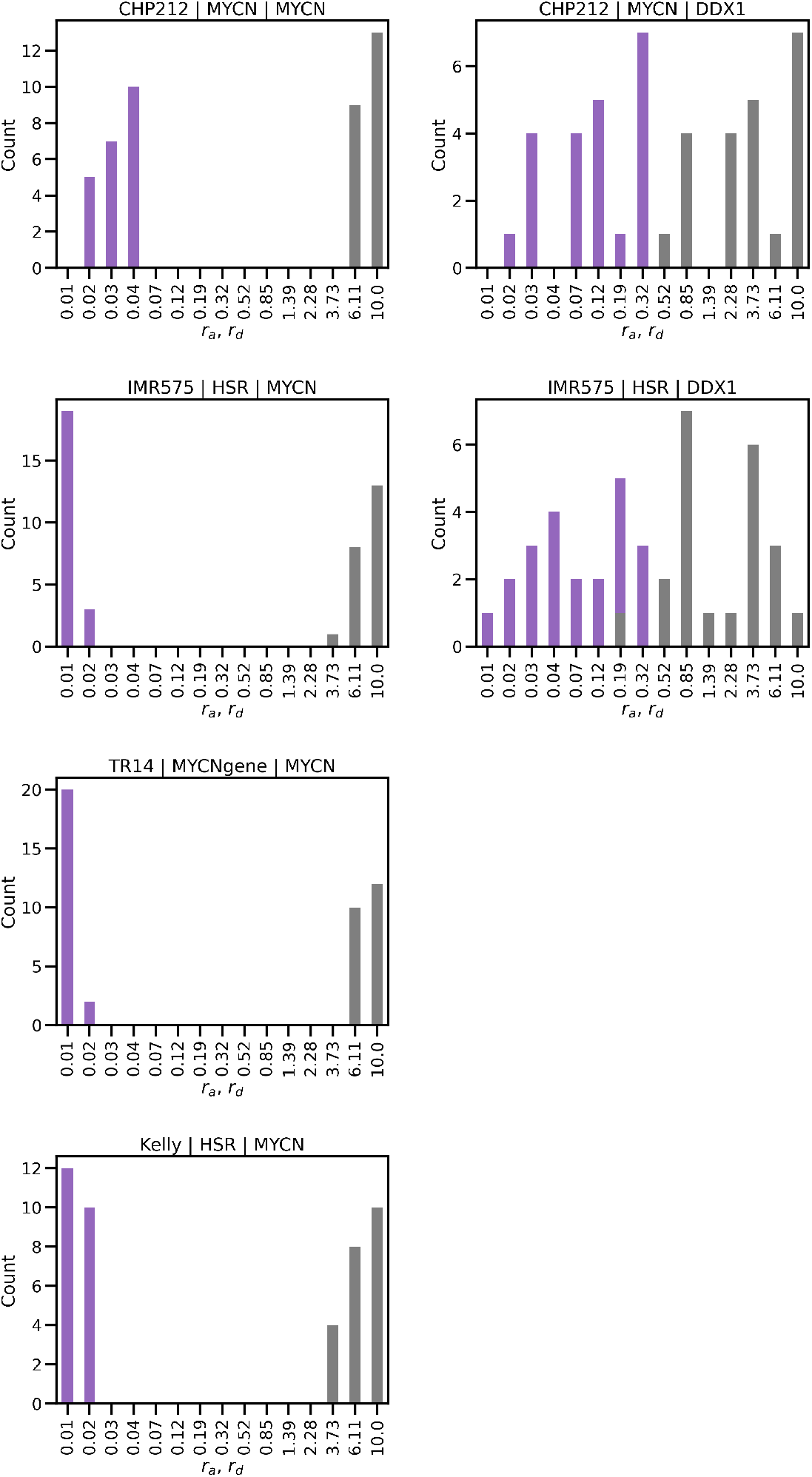
Posterior distributions for switching parameters obtained by fitting to the same gene (MYCN and DDX1) across different cell lines

It has been proposed that ecDNA form hubs, which contribute to the modulation and enhancement of cargo oncogenes via mechanisms such as enhancer sharing [15]. It has also been suggested that, in certain contexts, differences in copy number alone are important and that hub formation is not observed [16]. Our model fits seem to suggest a more nuanced explanation. On the one hand, the similarity of activation and deactivation rates indicates that at least the expression of some genes occurs independent of context, that copy number differences alone are sufficient to explain the heightened expression observed in ecDNA contexts. On the other hand, modulation of activation and deactivation rates is a much more plastic mechanism than variations in ecDNA copy number through progressive loss over time, and is moreover dependent on the presence of regulatory elements such as enhancers. Thus, in the presence of ecDNA hubs, these rates can be different and contribute to differentiated expression.

### 3.4 Genotoxic treatment is predicted to give rise to heterogeneous persister cell populations

We now discuss how plasticity arising from stochastic gene transcription impacts tumour progression, particularly treatment response. We simulated the effect of genotoxic therapy and investigated the impact of transcription stochasticity on treatment outcomes *in silico* (Figure 5). We observed that cells surviving treatment differ substantially across different parameter regimes, i.e. *r*_*a*_ *< r*_*d*_, *r*_*a*_ = *r*_*d*_ and *r*_*a*_ *> r*_*d*_.

**Figure 5.**
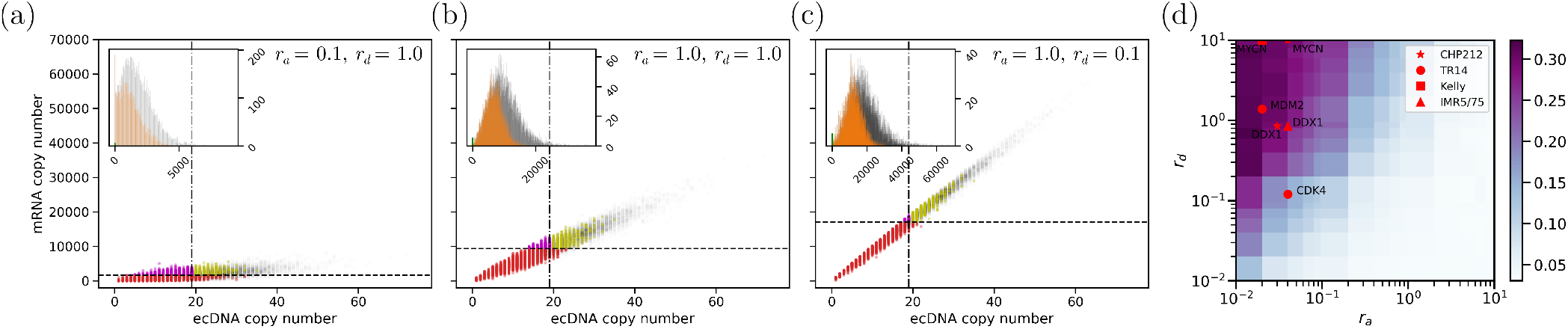
Joint and marginal distributions showing persisting cells after genotoxic treatment

First, treatment shifted the marginal mRNA distribution and its peak towards lower values across all parameter regimes before (inset panels grey) and after treatment (inset panels orange). Next, using joint ecDNA–mRNA distributions, we classified persisting cells after treatment into subpopulations depending on their mRNA content prior to treatment: ‘low-mRNA’ cells with lower than median mRNA copies (red dots); ‘low-ecDNA, high-mRNA’ cells with higher than median mRNA copies but lower than median ecDNA copies (pink dots); and ‘high-ecDNA, high-mRNA’ cells with higher than median mRNA copies and higher than median ecDNA copies (yellow dots). The ‘low-ecDNA, high-mRNA’ cells (pink dots) are the subpopulation that we expect to be most aggressive (mRNA copy number taken as proxy) and least amenable to further genotoxic treatment (because of low ecDNA copy number). The fraction of this subpopulation in surviving cells is highest when *r*_*a*_ *< r*_*d*_ (Figure 5a).

As shown previously, when *r*_*a*_ *> r*_*d*_, the shapes of the ecDNA and mRNA copy number distributions coincide. In this regime, a treatment targeting DNA copies functions effectively to reduce the mRNA copy numbers as well. However, when *r*_*a*_≤ *r*_*d*_, a treatment targeting DNA copies may not always reduce mRNA copy numbers as well. This should be considered when applying the same treatment in different oncogene contexts. Using our estimated *r*_*a*_ and *r*_*d*_ values for oncogenes in each cell line, we show the predicted fraction of surviving ‘low-ecDNA, high-mRNA’ cells for different genes (Figure 5d). Our predictions reveal a greater fraction of such cells in the case of *MYCN* and *MDM2* than in the case of *CDK4*.

## 4 Discussion

ecDNA are an important contributing factor to intratumour heterogeneity, both through copy number variation and gene expression heterogeneity. We used a two-step stochastic model of transcription to build a general framework to study gene expression on ecDNA. Our model recapitulated experimentally observed patterns in single-cell ecDNA–mRNA data, paving the way for a quantitative understanding of differences in expression between genes and amplification contexts.

Although co-amplification of multiple genes has been observed in the same ecDNA elements in various tumours [17, 18], their expression level can vary significantly [4, 19]. ecDNA copy numbers and mRNA have been observed to be positively correlated and previous studies have used linear fitting to quantify such correlations; however, the slope of such linear fits and the variance are different for different genes [4]. Our model provides a simple framework to quantify the relationship between DNA and mRNA copy number distributions beyond linear correlations. We can distinguish between genes through different inferred parameter ranges of gene activation and deactivation rates. For example, while *MYCN* is overexpressed in all cell lines, the best fit for *MYCN* in ecDNA cell lines is the combination of low activation and high deactivation rates, which is different from parameter estimates for other overexpressed genes such as *TRIB2, LPIN1* and *DDX1* in the same cell lines. This might explain why high copy numbers of *MYCN* are often observed in neuroblastoma tumours [13].

Interestingly, similar parameter ranges were inferred for the genes *MYCN* and *DDX1* in neuroblastoma cell lines with ecDNA (CHP212 and TR14) and those with HSR amplifications (Kelly, IMR5/75), suggesting that differences in expression in this context are primarily driven by differences in gene copy number. This touches on a key question in the field: whether expression on ecDNA is fundamentally different to expression on chromosomal DNA. According to some recent studies, ecDNA could have more accessible chromatin than chromosomal DNA (including HSR), leading to higher expression per copy in ecDNA, such as that observed for certain genes in TCGA samples [20]. Additionally, it has been proposed that ecDNA form ‘hubs’, where interactions between genes and enhancers lead to a higher level of expression compared to chromosomal DNA [15]. However, other studies have found that expression per copy number is unchanged between ecDNA and chromosomal DNA [16]. The picture that emerges is that expression across different types of amplification in cancer can be complex and patient- or context-dependent. While in this study we were limited by the available single-cell, matched ecDNA–mRNA data, our quantitative method can be extended and applied to other oncogenes in cell lines or patients using bulk or single-cell data.

The variation among mean ecDNA copy numbers among different patients with the same tumour types and ecDNA oncogenes indicates that the expression level per copy number might have large intratumour heterogeneity. Our model of stochastic gene expression showed that varying the activation and deactivation rates of oncogenes does not only change the effective expression per copy number, but also has a strong impact on how much individual cells with the same or similar gene copy number vary in their overall expression. Changing these rates is thus a mechanism for introducing variation in shorter timescales compared with changes in ecDNA copy numbers through random segregation during cell division and selection.

Using our model to understand the effect of transcriptional heterogeneity on treatment response is another avenue for future work. Simulating genotoxic therapy, we predicted that different fractions of cells with high mRNA and low ecDNA content would survive treatment depending on the activation and deactivation rates of the focused genes, leading to variable susceptibility to further treatment.

In the present study we used a minimal, two-step model of transcription to study the effect of stochasticity. While this model was able to capture patterns in experimental distributions, it does not take into account interactions between different genes that are observed to be co-amplified in cancer. Extending our model to include gene–gene interactions and the effect of a growing population would allow us to model phenotypic heterogeneity with greater accuracy.

