## Supplementary materials for "Mathematical Modelling of Stochastic Gene Expression Quantifies mRNA Heterogeneity in Cancer Cell Populations with Extrachromosomal DNA"

### Supplementary figures

##### IMR5/75

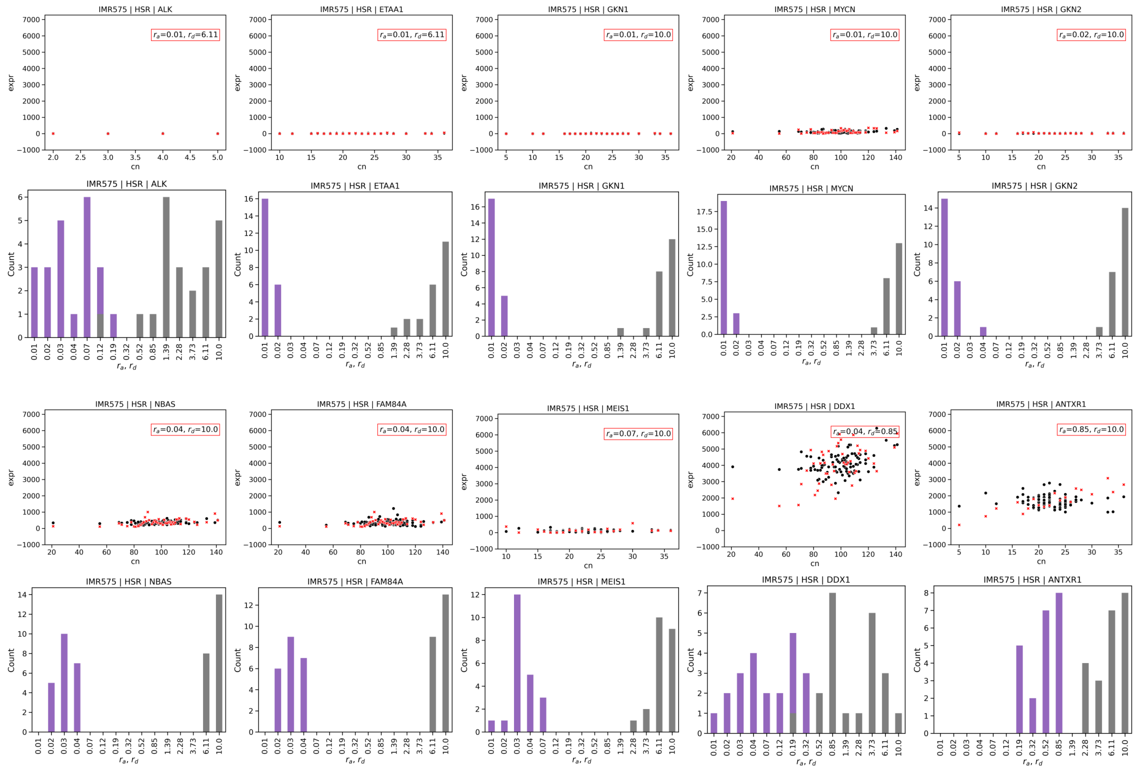


##### Kelly

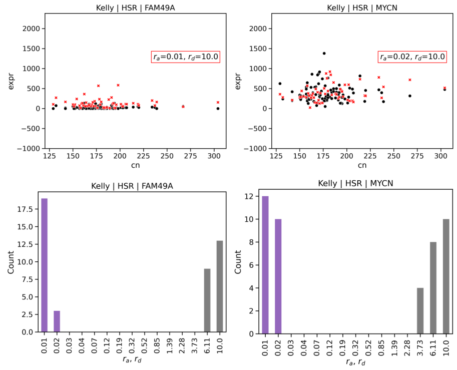


Figure S1: Posterior distributions and best fits for all genes in IMR5/75 and Kelly cell lines.

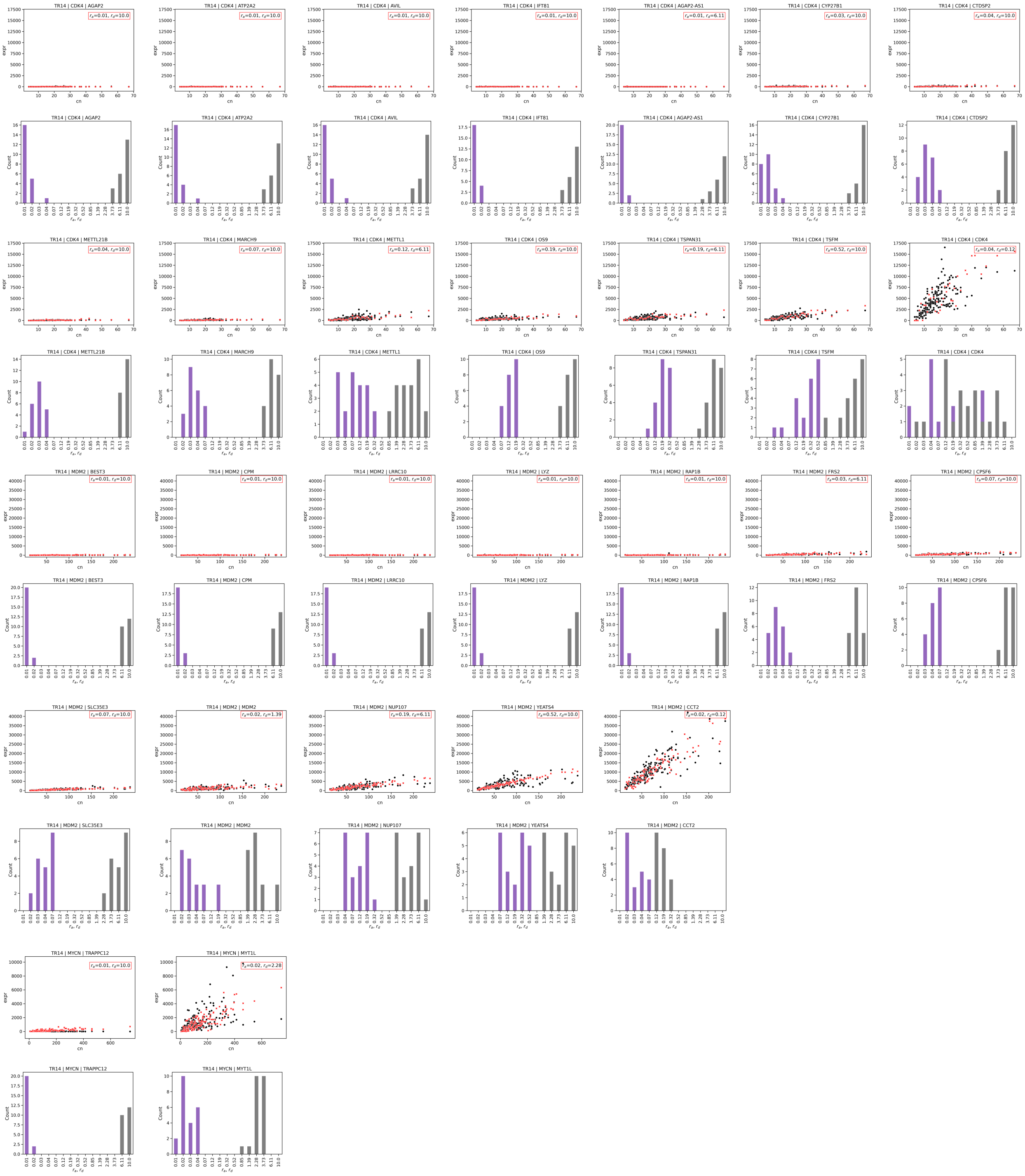


Figure S2: Posterior distributions and best fits for all genes in the TR14 cell line.

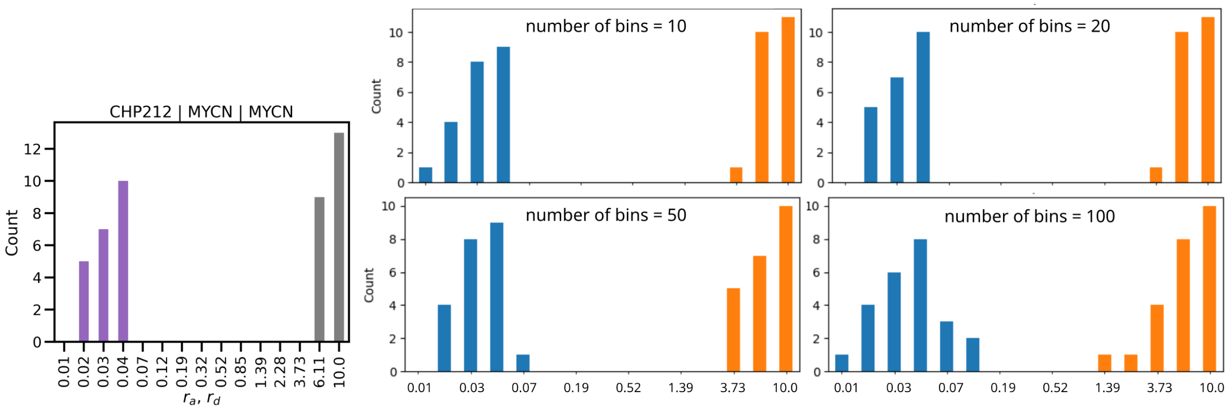


Figure S3: Posterior distribution estimated from joint distribution (leftmost panel) and posteriors estimated from marginal distributions using different numbers of bins (four right panels, $r_a$ shown in blue, $r_d$ shown in orange).

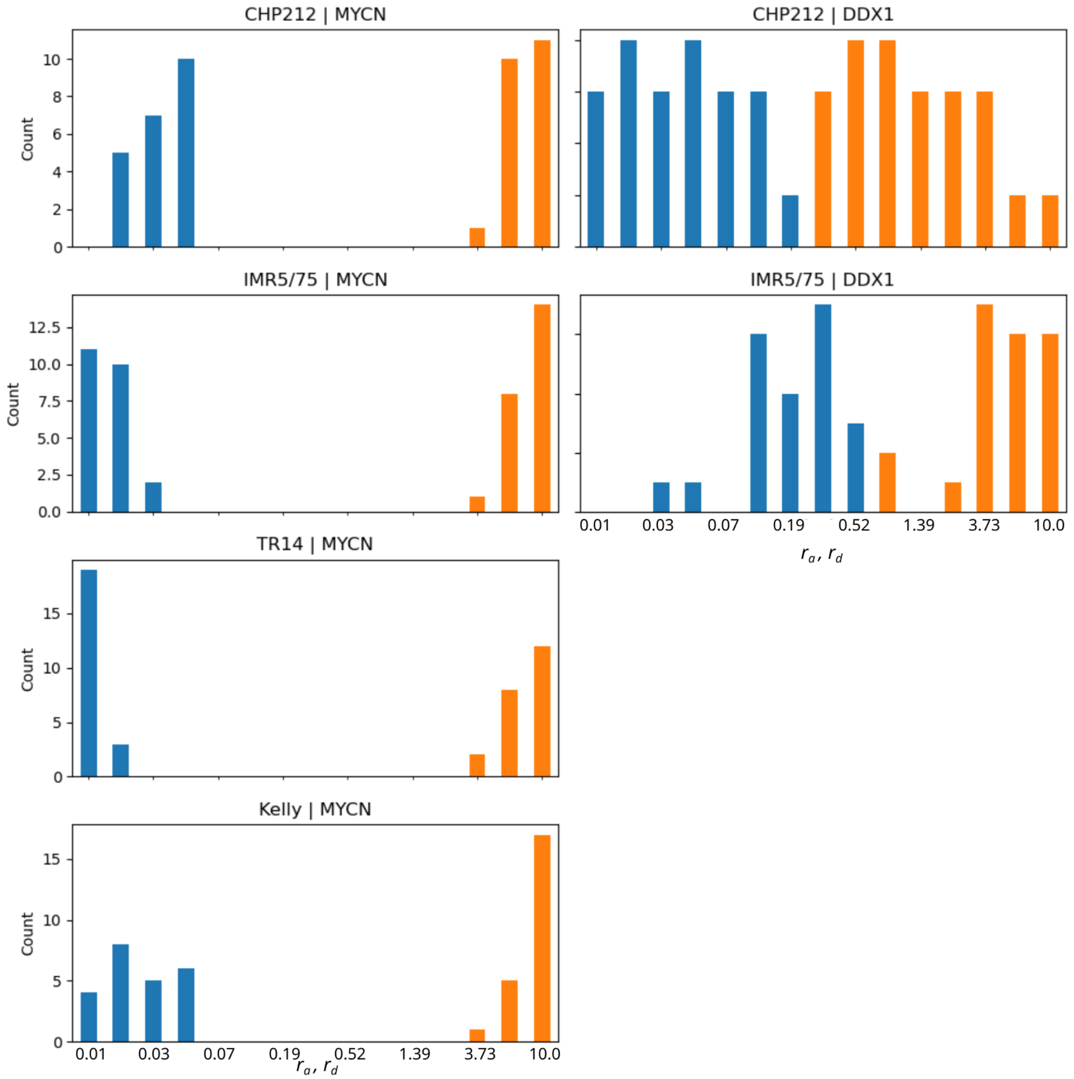


Figure S4: Posterior distributions for *MYCN* and *DDX1* obtained using marginal distributions (number of bins=20).

#### Steady-state solution for isogenic populations with one gene copy per cell

From Karmakar and Bose [?], the probability density of obtaining an mRNA concentration of $x$ ($x \in [0,1]$) at time $t$ is a sum of two terms, each corresponding to a gene state ("off" or "on"):

$$p(x,t) = p_0(x,t) + p_1(x,t). \tag{1}$$

Gene switching is assumed to be the only source of stochasticity in the system, with mRNA being produced and decaying deterministically according to the equation

$$\frac{dx}{dt} = \frac{j_m}{X_{max}}\, z - k_m\, x =: f(x,z), \tag{2}$$

where $X_{max}$ is the maximum concentration of mRNA, $z \in \{0,1\}$ is the gene state, $j_m$ is the rate of mRNA production and $k_m$ is the rate of mRNA decay.

The probability densities $p_z(x,t), z \in \{0,1\}$ change according to the equation:

$$\frac{\partial p_z(x,t)}{\partial t} = -\frac{\partial}{\partial x}[f(x,z)p_z(x,t)] + \sum_{k \neq z}[W_{kz}\, p_k(x,t) - W_{zk}\, p_z(x,t)], \tag{3}$$

where $W$ is the matrix of transition rates. Thus, the time derivatives of densities $p_0, p_1$ are given by:

$$\frac{\partial p_0(x,t)}{\partial t} = -\frac{\partial}{\partial x}[-k_m\, x\, p_0(x,t)] + [k_d\, p_1(x,t) - k_a\, p_0(x,t)] \tag{4}$$

$$\frac{\partial p_1(x,t)}{\partial t} = -\frac{\partial}{\partial x}\Big[\Big(\frac{j_m}{X_{max}} - k_m\, x\Big)p_1(x,t)\Big] + [k_a\, p_0(x,t) - k_d\, p_1(x,t)]. \tag{5}$$

At steady state: $\frac{\partial p_z}{\partial t} = 0, z = \{0,1\}$, which implies

$$-\frac{d}{dx}[-x\, p_0^{st}(x)] + r_d\, p_1^{st}(x) - r_a\, p_0^{st}(x) = 0 \tag{6}$$

$$-\frac{d}{dx}[(1-x)p_1^{st}(x)] + r_a\, p_0^{st}(x) - r_d\, p_1^{st}(x) = 0, \tag{7}$$

where $r_a = k_a/k_m$ and $r_d = k_d/k_m$, and we have used $X_{max} = j_m/k_m$. Adding the two ODEs above we get:

$$\frac{d}{dx}\Big(x\, p_0^{st} + (x-1)\, p_1^{st}\Big) = 0 \tag{8}$$

which implies $x\, p_0^{st} + (x-1)\, p_1^{st}$ is a constant. Substituting $x = 0$ and using the boundary condition $p_1^{st}(0) = 0$, we determine the constant to be equal to 0. Thus,

$$p_1^{st}(x) = \frac{x}{1-x}\, p_0^{st}(x). \tag{9}$$

Substituting for $p_1^{st}$ in (6) we get:

$$\frac{d}{dx}[x\, p_0^{st}(x)] + \Big(r_d\, \frac{x}{1-x} - r_a\Big)p_0^{st}(x) = 0. \tag{10}$$

In terms of the total probability density, $p_0^{st}(x) = (1-x)p^{st}(x)$, which means

$$\frac{d}{dx}[x\,(1-x)\, p^{st}(x)] + \Big(r_d\, x - r_a\,(1-x)\Big)p^{st}(x) = 0. \tag{11}$$

Defining $f(x) := x\,(1-x)\, p^{st}(x)$, it follows that

$$f'(x) + \Big(\frac{r_d}{1-x} - \frac{r_a}{x}\Big)f(x) = 0, \tag{12}$$

where $f'(x) = \dfrac{df(x)}{dx}$. Integrating over $x$, the solution to the above equation is found to be

$$f(x) = C\, x^{r_a}\, (1-x)^{r_d}, \tag{13}$$

which gives us the steady state solution

$$p^{st}(x) = C\, x^{r_a - 1}\, (1-x)^{r_d - 1}. \tag{14}$$

Because $\int_0^1 p^{st}(x) dx = 1$, the normalisation constant is $C = \dfrac{1}{\int_0^1 x^{r_a-1}\,(1-x)^{r_d-1} dx} = \dfrac{1}{B(r_a, r_d)}$, where $B(r_a, r_d)$ is the beta function.

### Generalisation to more than one gene copy

Let's assume we have two copies of the gene of interest in each cell. We also assume independent activation and deactivation of the copies. The steady-state ODEs of the system are given by:

$$-\frac{d}{dx}[-x\, p^{st}_{0,0}(x)] + r_d\, p^{st}_{1,0}(x) - r_a\, p^{st}_{0,0}(x) + r_d\, p^{st}_{0,1}(x) - r_a\, p^{st}_{0,0}(x) = 0 \tag{15}$$

$$-\frac{d}{dx}[(1/2 - x)\, p^{st}_{1,0}(x)] + r_a\, p^{st}_{0,0}(x) - r_d\, p^{st}_{1,0}(x) + r_d\, p^{st}_{1,1}(x) - r_a\, p^{st}_{1,0}(x) = 0 \tag{16}$$

$$-\frac{d}{dx}[(1/2 - x)\, p^{st}_{0,1}(x)] + r_d\, p^{st}_{1,1}(x) - r_a\, p^{st}_{0,1}(x) + r_a\, p^{st}_{0,0}(x) - r_d\, p^{st}_{0,1}(x) = 0 \tag{17}$$

$$-\frac{d}{dx}[(1 - x)\, p^{st}_{1,1}(x)] + r_a\, p^{st}_{0,1}(x) - r_d\, p^{st}_{1,1}(x) + r_a\, p^{st}_{1,0}(x) - r_d\, p^{st}_{1,1}(x) = 0 \tag{18}$$

Changing the notation slightly so that the subscript of the probability density function $p$ now denotes the *number of active genes* and not the gene state, that is:

$$p_0 := p_{0,0}, \quad p_1 := p_{0,1} + p_{1,0}, \quad p_2 := p_{1,1},$$

and assuming the superscript and $x$ dependence to be implied, we can write the following equations to describe the steady state of the system:

$$-\frac{d}{dx}[-x\, p_0] + r_d\, p_1 - 2\, r_a\, p_0 = 0 \tag{19}$$

$$-\frac{d}{dx}[(1/2 - x)\, p_1] + 2\, r_a\, p_0 + 2\, r_d\, p_2 - (r_a + r_d) p_1 = 0 \tag{20}$$

$$-\frac{d}{dx}[(1 - x)\, p_2] + r_a\, p_1 - 2\, r_d\, p_2 = 0. \tag{21}$$

Using these equations, we can express $p_1$ in terms of $p_0$ and $p_2$ as:

$$p_1 = \frac{x}{1/2 - x} p_0 - \frac{1 - x}{1/2 - x} p_2. \tag{22}$$

The system of ODEs can be reduced to two coupled equations in terms of $p_0$ and $p_2$:

$$p_0' = f(x, r_a, r_d)\, p_0 + g(x, r_d)\, p_2 \tag{23}$$

$$p_2' = h(x, r_a)\, p_0 + j(x, r_a, r_d)\, p_2, \tag{24}$$

where

$$f(x, r_a, r_d) = \frac{2r_a - 1}{x} + \frac{r_d}{x - 1/2} \tag{25}$$

$$g(x, r_d) = \frac{r_d\, x}{x\,(x - 1/2)} \tag{26}$$

$$h(x, r_a) = \frac{r_a\, x}{(x-1)\,(x - 1/2)} \tag{27}$$

$$j(x, r_a, r_d) = \frac{2r_d - 1}{x - 1} + \frac{r_a}{x - 1/2}. \tag{28}$$

We can solve the two coupled ODEs exactly for special cases: $r_a = 0$, $r_d = 0$. For general nonzero values of $r_a, r_d$ it is challenging to find an analytical solution for $p_j, j = 0, 1, 2$.

Instead of finding an exact solution for the steady-state probability density function, we ask how many peaks the pdf has as a function of $x$. At fixed points in $x$, $p_0'(x) = 0, p_2'(x) = 0$. Thus, we look for soluions to the following coupled equations:

$$f(x, r_a, r_d)\, p_0 + g(x, r_d)\, p_2 = 0 \tag{29}$$

$$h(x, r_a)\, p_0 + j(x, r_a, r_d)\, p_2 = 0. \tag{30}$$

Multiplying the first equation by $h(x, r_a)$, the second equation by $f(x, r_a, r_d)$ and subtracting the two resultant equations, we get

$$[g(x, r_d), h(x, r_a) - j(x, r_a, r_d)\, f(x, r_a, r_d)]p_2 = 0. \tag{31}$$

For $r_a = r_d$, we can solve the resulting quadratic equation in $x$ using symbolic computation. The equation has the solutions

$$x = \left\{ \frac{1 - 6r_a + 9r_a^2 - \sqrt{2r_a^2 - 12r_a^3 + 17r_a^4}}{2(1 - 6r_a + 8r_a^2)}, \frac{1 - 6r_a + 9r_a^2 + \sqrt{2r_a^2 - 12r_a^3 + 17r_a^4}}{2(1 - 6r_a + 8r_a^2)} \right\}$$

The denominator of the solution blows up for $r_a = \frac{1}{4}, \frac{1}{2}$. Consequently, the solutions change behaviour at these points.

Using this intuition from two gene copies, we conjecture that for $n$ gene copies, the behaviour of the system will differ for $r_a, r_d < 1/n$ (slow switching) and $r_a, r_d > 1/n$ (fast switching). We use this to determine parameter sweep regions for numerical simulations in the main text.
